# Hippocampal-striatal functional connectivity supports processing of temporal expectations from associative memory

**DOI:** 10.1101/699439

**Authors:** Vincent van de Ven, Chanju Lee, Julia Lifanov, Sarach Kochs, Henk Jansma, Peter De Weerd

## Abstract

The hippocampus and dorsal striatum are both associated with temporal processing, but they are thought to play distinct roles. The hippocampus has been reported to contribute to storing temporal structure of events in memory, whereas the striatum contributes to temporal motor preparation and reward anticipation. Here, we asked whether the striatum cooperates with the hippocampus in processing the temporal context of memorized visual associations. In our task, participants were trained to implicitly form temporal expectations for one of two possible time intervals associated to specific cue-target associations, and subsequently were scanned using 7T functional magnetic resonance imaging. During scanning, learned temporal expectations could be violated when the pairs were presented at either the learned or not-learned time intervals. When temporal expectations were not met during testing trials, activity in hippocampal subfields CA3/CA2 and CA1 decreased while right putamen activity increased, compared to when temporal expectations were met. Further, psycho-physiological interactions showed that functional connectivity between left CA1 and caudate, as well as between putamen and caudate, decreased when temporal expectations were not met. Our results indicate that the hippocampus and striatum cooperate to process implicit temporal expectation from mnemonic associations, with different but complementary contributions from caudate and putamen. Our findings provide further support for a hippocampal-striatal network in temporal associative processing.

## Introduction

Extensive research on episodic memory has supported the suggestion that the ability to correctly order events into a coherent and continuous sequence (Tulving, 1984; Kurby and Zacks, 2008) is crucial for various cognitive abilities and functioning of our daily life (*e.g.*, (Vargha-Khadem et al., 1997; Bartsch et al., 2011; Baker et al., 2014)). There is rapidly growing consensus from both animal neurophysiological and human neuroimaging research that the medial temporal lobe, including the hippocampus, is involved in representing temporal information in memory (Eichenbaum, 2013, 2014; Ranganath and Hsieh, 2016). Yet, processing of temporal information has long been associated with activity in the dorsal striatum and other parts of the motor circuit (Matell et al., 2003; Meck et al., 2008; Mello et al., 2015). Further, the dorsal striatum and hippocampus can show increased functional connectivity during memory encoding or retrieval in spatial associative contexts. Investigation of hippocampal-striatal interaction during temporal associative contexts has not yet been described. This was the aim of the current study.

The medial temporal lobe, specifically the hippocampus, has been suggested as a primary brain region for processing spatial navigation and episodic memory (Squire, 1992; Milner et al., 1998; Bird and Burgess, 2008). Over the last decade, researchers reported that time is also processed in the hippocampus (Eichenbaum, 2014). Single cell recordings in the rat hippocampus found peak firing of hippocampal cells at successive moments during delay periods inserted between cue and probe within trials of a paired associate task (Pastalkova et al., 2008; MacDonald et al., 2011, 2013). The activities of these so-called “time cells” might reflect encoding of the temporal dimensions of events, a crucial property of episodic memory (Tulving, 1984; Ergorul and Eichenbaum, 2004). This argument was further supported by lesion studies with rats that showed hippocampal damage results in disruption of memory for time without impairing the recognition of items in learned sequences (Ergorul and Eichenbaum, 2004; DeVito and Eichenbaum, 2011).

Similar to rodent research, human neuroimaging research also observed activity in the hippocampus that is compatible with temporal associative memory. A number of studies discovered differential activations in the hippocampus during tasks in which temporal contexts changed (*e.g.*, (Staresina and Davachi, 2009; Schapiro et al., 2012; Ezzyat and Davachi, 2014)). Importantly, the hippocampus seemed to be mostly related to bridging the temporal gap between objects that were presented sequentially and in close temporal proximity. For example, one study investigated the effects of temporal order of memorized sequences of objects (Ezzyat and Davachi, 2014), and found that hippocampal patterns were more similar for objects when their positions were temporally close within a learned sequence. Thus, rather than a clock, the hippocampus might function as an associator of information across different moments in time. In further support of this notion are findings that suggest that the hippocampus may encode relative temporal structure at different time scales (Mankin et al., 2015).

In parallel lines of research, the dorsal striatum (DS), including caudate nucleus and putamen, has long been strongly associated with temporal processing of stimulus-response events (Buhusi and Meck, 2005; Coull et al., 2010; van Rijn et al., 2014). Neurophysiological DS cells have been found to increase activity when a behaviorally relevant period of time is about to be over (Matell et al., 2003; Mello et al., 2015). Further, DS lesions in rats (Meck, 2006), neurological diseases affecting dorsal striatal areas (Malapani et al., 1998), and disruptions in DS dopamine signaling (Rowe et al., 2010), interfere with the processing of time. In addition to the findings from rodent research, human neuroimaging studies also reported higher striatal activity when participants were engaged in tasks that required the processing of interval timings (Rao et al., 2001; Ferrandez et al., 2003; Coull et al., 2004; Tanaka et al., 2004). The majority of these studies focused on motor preparation to timed or rhythmic responses, with little to no investigation of the possible relation to episodic memory formation. However, DS, particularly putamen, may also be involved in processing violations of temporal expectancy about reward delivery (Mcclure et al., 2003; Doherty et al., 2004; Seymour et al., 2004). Mcclure et al. (Mcclure et al., 2003) used a classical conditioning paradigm in which participants either learned that a reward was delivered six seconds after cue onset or after an unpredictable interval. During the test phase, the reward in the previously predictable context could now on some trials unexpectedly be delivered four seconds later. Results showed increased left putamen activity for the unexpectedly delayed reward, compared to reward delivered at the predicted interval of six seconds, suggesting that DS processes temporal prediction errors that indicate violation of expectations.

Interestingly, the hippocampus has shown increased functional connectivity with DS during encoding of new episodic (Sadeh et al., 2011) or associative memories (Mattfeld and Stark, 2015). Moreover, both subcortical structures appear to be involved in spatial processing and navigation through a real or virtual environment (Voermans et al., 2004; Gengler et al., 2005; Igloi et al., 2010), suggesting that they interact when processing the contextual aspects of associative events in memory encoding. Whether this interaction also plays a role in associative memory of time has not yet been investigated.

The purpose of this study was to test whether associative temporal memory is related to hippocampal-striatal connectivity in the human brain. To this end, we had participants learn cue-target associative pairs of visual stimuli in different temporal contexts, in the form of different time intervals between a cue and target (van de Ven et al., 2017). An important property of these memories was that each cue was hypothesized to elicit neural responses representing the prediction of the temporal context in which the associated target event would follow in the near future, *i.e.*, *when* the associated target would appear. During memory testing, participants could be presented with cue-target pairs in the learned as well as in a novel temporal context. This discrepancy of brain activity in different temporal contexts was measured using ultra-high field (UHF) functional magnetic resonance imaging (fMRI) at 7 Tesla. The main analyses focused on dorsal striatal structures and hippocampal subfields in the left and right hemispheres, in which we analyzed regional activity and inter-regional connectivity as a function of memory-based temporal expectancy. We hypothesized that striatum and hippocampus would show higher activity when temporal expectancies from memory were not met, thereby signaling increased temporal prediction error. Further, we hypothesized decreased hippocampal-striatal functional connectivity, as analyzed using psychophysiological interactions (PPI) when temporal expectancies were not met.

## Methods

### Participants

Eighteen healthy young adults (mean [SD] age = 23.06 [3.02] years, 15 females) participated in the study. To ensure suitability with the MR environment, all participants were screened by experimenters before participation. All participants provided written informed consent to participate in the study and MR measurements, and received financial compensation for their participation. The study was approved by the local ethics committee of the Faculty of Psychology and Neuroscience (FPN) of Maastricht University.

### Task procedure

Participants completed a time paired associative task (van de Ven et al., 2017) in which they learned to associate pairs of cue-target stimuli, which were separated by one of two time delays. For the stimuli, we used eight pairs of abstract shapes to minimize conceptual processing and make the task challenging. Associated stimulus pairs were randomly created for each participant.

Participants first learned the cue-target pairs outside the MR scanner, with the task presented on a laptop screen. During the testing phase, stimuli were delivered to participants at the same visual angles through a mirror system while lying in the MR scanner. Each stimulus was shown at a size of 5°×5° visual angle at the center of the screen, on a grey surface. When no stimulus was presented, a fixation cross was presented at a size of 1°×1° visual angle. The experiment was programmed in Psychopy version 1.8 (Peirce, 2007), using its feature of screen refresh readout (refresh rate = 60 Hz) to maximally control stimulus and interval timing (Garaizar and Vadillo, 2014).

The task started with a learning phase (see **Figure 1**). At the start of the learning phase, the eight cue-target pairs were shown to participants once and without the requirement to respond to the items (passive exposure trials). Each trial began with the presentation of a fixation cross (500 msec) after which the cue item was shown (1000 msec). After cue offset, the target item was shown for 1000 msec at a delay interval of either 500 or 2000 msec (respectively L1 and L2 for short or long intervals during learning), with each pair assigned to one of the two intervals. Participants were not informed about the different temporal contexts. Participants then trained to learn and memorize the eight cue-target pairs. Each trial was similar in design as for the passive exposure trials, with two important exceptions. First, the latter item (referred to as probe) of each presented pair could either be the associated target (as seen during the passive exposure trials) or one of the seven non-target alternatives, randomly drawn on each non-target trial. Second, probes that were targets were always shown after the associated interval. When probes were non-targets, the interval was always the non-associated delay. Cues were shown with the associated targets (and thus with their associated intervals) in 50% of the trials. Participants had to determine whether the probe item was the cue-associated target and indicated their decision through a button press response within 3000 ms after probe onset. Response feedback (color change of the fixation cross indicating a correct [green] or incorrect [red] response) was provided after each trial to facilitate learning of the stimulus associations. One learning block consisted of 32 trials, which were presented in random order, and participants repeated learning blocks until reaching 84% accuracy within a learning block or until a maximum of 6 learning blocks were completed.

**Figure 1.**
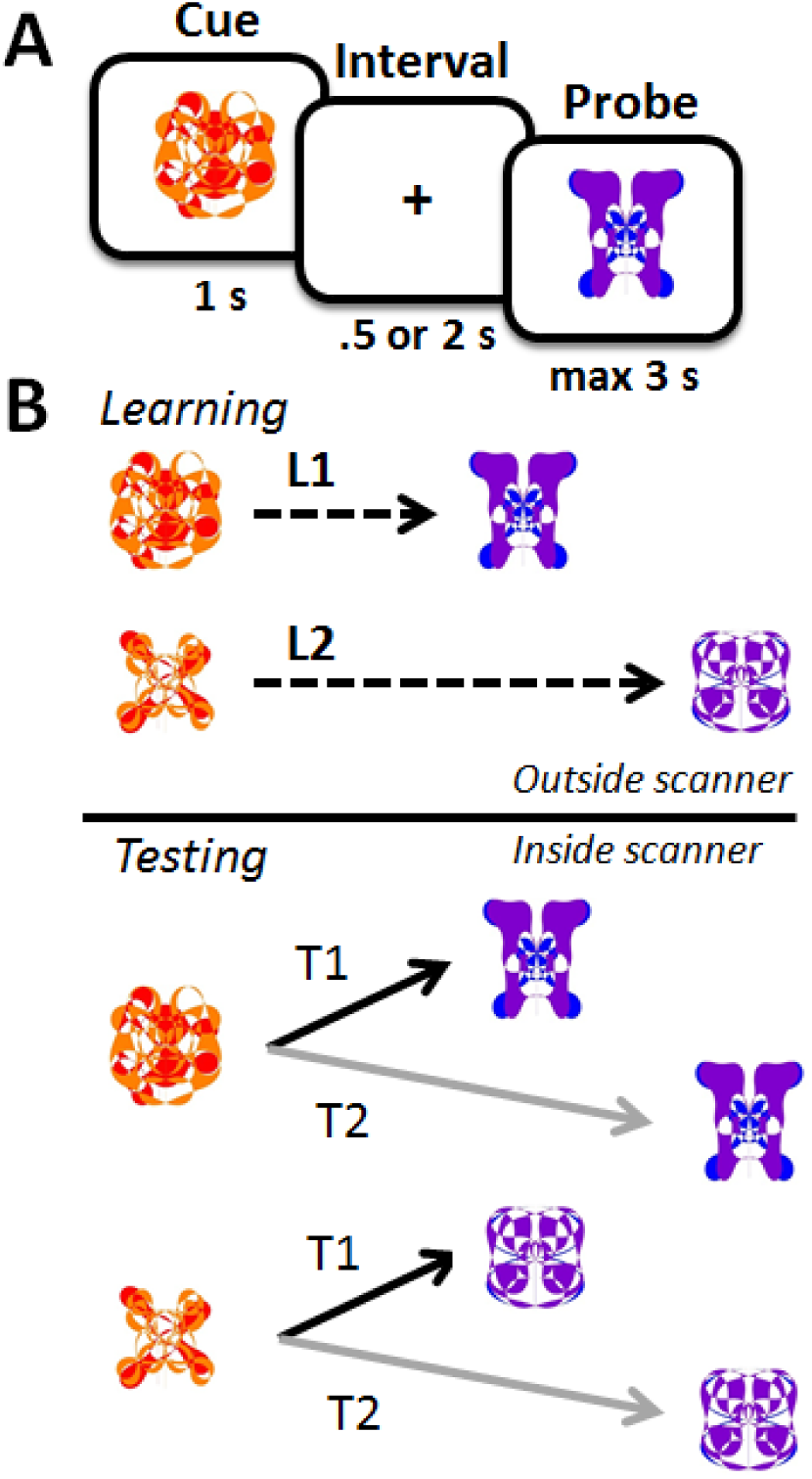
Task design. Participants had to learn cue-target associations, in which items were presented sequentially. Each pair was further association with one of two time intervals, L1 (500 ms) or L2 (2000 ms). During memory testing, each cue-target pair could be shown with either of the two intervals, T1 (500 ms) and T2 (2000 ms).

After the learning phase, participants underwent the testing phase in the MR scanner. The testing phase was similar to the learning phase, with two crucial differences. First, the pairing between cue and interval was broken, such that each cue was shown with either of the two intervals with equal probability of 0.5, regardless of whether the probe matched the target or not. We denote intervals between the cue and probe in the testing phase as T1 (500 ms) and T2 (2000 ms). Second, participants were not given any feedback about their responses throughout the testing phase. Trial order was randomized for each block and each participant. One testing block consisted of 64 trials and participants completed 2-3 testing blocks in the scanner. The inter-trial interval for the testing phase was jittered around an average of 8000 ms. After the session was completed, participants were asked whether they noticed any changes in temporal gaps between cue-target stimuli. An entire testing session (inside and outside of the scanner) lasted approximately 90-120 minutes and concluded with debriefing.

### MRI data acquisition

A Siemens MAGNETOM 7 Tesla MR scanner with a 32-channel head coil was used to acquire whole-brain imaging data. An EPI sequence was used to collect blood oxygenation level-dependent (BOLD) images. All scanning sessions were held at the Scannexus facility in Maastricht, the Netherlands. Participants were instructed to fixate their head and posture throughout the scanning. Before collecting any images, semi-automated shimming was performed. After the shimming, anatomical data were acquired using T1-weighted and Proton density-weighted images (0.7mm isotropic, 240 slices, no inter-slice gap, TR=5s). Functional images were collected using T2*-weighted images (1.25mm isotropic, 60 slices, no inter-slice gap, TR=1.5s, TE=22ms, FA=50, anterior-to-posterior phase direction). Speed of data acquisition was increased using a multi-band acquisition sequence of 2 simultaneous slices and a GRAPPA acceleration factor of 2 (Moeller et al., 2010). Additionally, five phase-inverted (posterior-to-anterior) EPI images were collected with the same imaging parameters for offline geometric distortion correction (see below). While participants were lying inside the scanner, experimental stimuli were delivered through a mirror system. During the functional runs, behavioral responses were simultaneously collected using an MR-compatible button box.

### Analysis

#### Preprocessing

Imaging data were preprocessed and analyzed using BrainVoyager v20.4 (Goebel et al., 2006) and the NeuroElf toolbox (http:/neuroelf.net) in MATLAB 2015a (www.mathworks.com). First, anatomical MR images were corrected for intensity inhomogeneity, skull-stripped and then spatially normalized to the MNI (Montreal Neurological Institute)-152 template space using an affine registration with 12 degrees of freedom. For functional images, processing steps included slice scan time correction (sinc interpolation), three-dimensional (3D) motion correction and temporal filtering using linear trend removal and high-pass filtering (4 sine/cosine cycles across the full timecourse). Geometrical distortions in functional images that resulted from EPI sequences at 7 Tesla were corrected using a set of five phase-inversed EPI baseline images with the Correction based on Opposite Encoding (COPE) plugin version 1.0, which follows a previously published offline image correction approach (Andersson and Skare, 2002; Andersson et al., 2003). Geometrically corrected and preprocessed functional images were then normalized to MNI space.

#### Region-of-interest (ROI) creation

For the purpose of this study, the hippocampus and striatum were *a priori* selected as regions-of-interests (ROIs). Bilateral hippocampus and its subfields were segmented from the T1 images of each participant individually using the online anatomical processing pipeline volBrain (http://volbrain.upv.es; (Manjón and Coupé, 2016)). In this pipeline, hippocampus and its subfields are localized and extracted from each anatomical image using a patch-based segmentation method (Coupé et al., 2011), which resulted in four ROIs for hippocampal subfield dentate gyrus/CA4 (DG/CA4), CA3/CA2, CA1 and Subiculum (Sub) in each hemisphere for each participant. Left and right striatal ROIs were taken from an anatomical atlas of the basal ganglia that was based on high-resolution 7T multi-modal MR images in young adults (Keuken et al., 2014). We separated the dorsal striatal ROIs in each hemisphere into the caudate nucleus and putamen using NeuroElf and BrainVoyager’s segmentation tools. **Figure 2** depicts the hippocampal and striatal ROIs used in the study.

**Figure 2.**
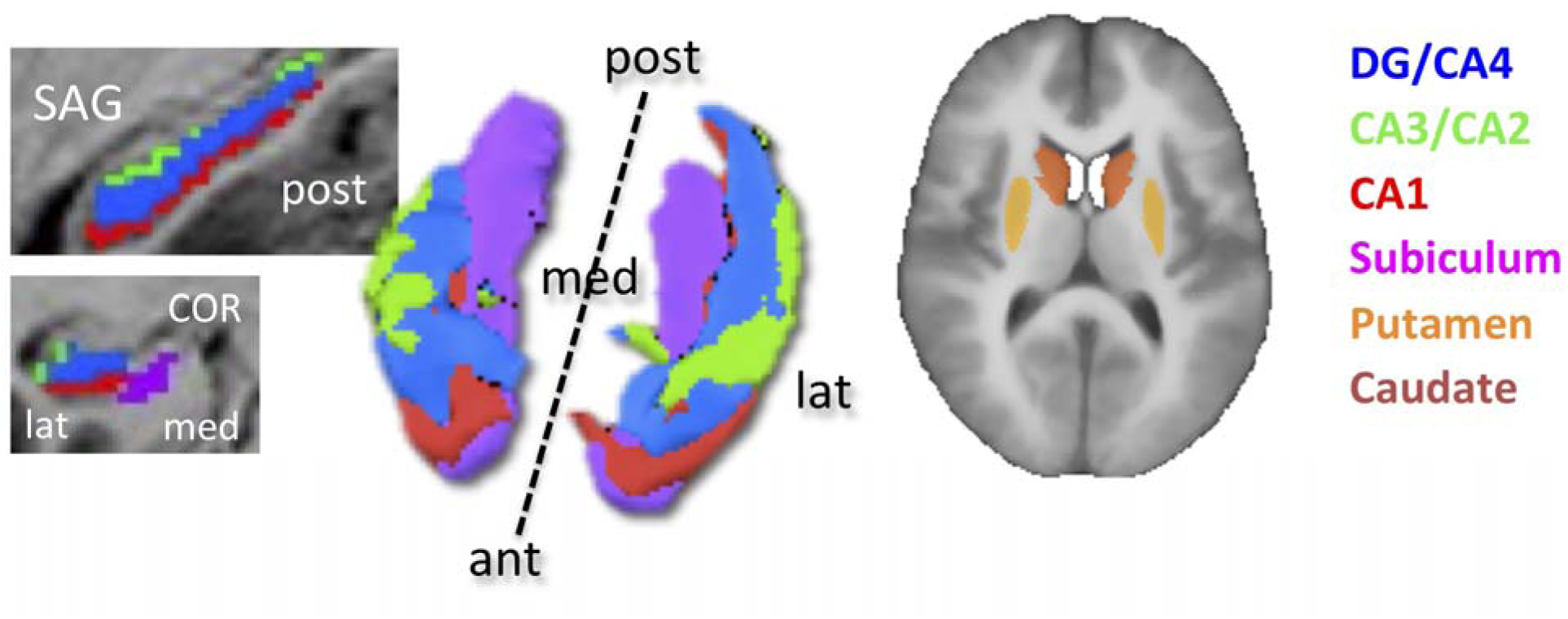
Hippocampal and dorsal striatal regions of interest (ROIs). ROIs are superimposed on the average anatomy of all participants. Color coding is shown in the figure. * P < 0.05, uncorrected; ** P < 0.05 Bonferroni-Holm corrected. Left hemisphere (marked by “L”) shown on the right in each panel.

#### Statistical analysis

Functional data were analysed using a multi-level general linear model (GLM) with the four interval-based task conditions (L1xT1, L1xT2, L2xT1 and L2xT2) containing correctly responded trials only and two additional conditions for inaccurately responded trials or trials with missing responses. GLM predictors were convolved with a two-gamma hemodynamic response function (HRF). The task regressors were appended with six (Z-normalized) head motion displacement vectors, as estimated by BrainVoyager’s head motion correction procedure (Goebel et al., 2006; Christoffels et al., 2011), and their first derivates. The GLM was applied at the ROI level, at which timeseries of voxels within an ROI were sampled and averaged for each participant. At the first analysis level, the GLM was fit to the functional data of each participant. At the second level, the single-subject GLM coefficients for the task conditions were analysed at the subject-level using a Random Effects (RFX) approach. Particularly, we were mainly interested in long-interval trials (that is, L1xT2 and L2xT2) as in these trials the interval was captured by at least one MR functional volume. Thus, we focused on the contrast [L1xT2 – L2xT2]. Statistical results were corrected for multiple comparisons (eight hippocampal and four striatal ROIs) using a false-discovery rate (FDR) of q=0.05 (Benjamini and Hochberg, 1995; Genovese et al., 2002).

In addition, an explorative voxel-by-voxel analysis was performed at the whole-brain level using the RFX GLM to explore activations that were induced by the task paradigm outside of the ROIs. Multiple comparison correction was performed at the cluster-level, using a Monte Carlo simulation of 1000 random statistical images of which values were drawn from a normal distribution and in which the spatial smoothness of each simulation was based on the empirical statistical map (Forman et al., 1995; Goebel et al., 2006). Clusters from the simulated maps were tabulated and ranked in size, from which the cluster size at a false positive rate of .05 was taken as minimum cluster threshold for visualizing the empirical map.

#### Functional connectivity analysis: psychophysiological interactions

Functional connectivity between ROIs engaged in our task paradigm was investigated using psychophysiological interactions (PPI) analysis (Friston et al., 1997; O’Reilly et al., 2012). The PPI design matrix, including the interaction term, was generated for each participant separately using the NeuroElf toolbox. For each PPI, the psychological and physiological variables were deconvolved prior to calculating the interaction term, which was then convolved with a two-gamma HRF. The PPI model was then tested in a similar two-level RFX approach as the task-based GLM, with regression fits estimated for each individual (first-level) combined into a group-level test of significance (second-level).

## Results

### Behavioral results

For analysis of the behavioral results, data of two participants who failed to press response buttons within the maximum response duration on more than half of the trials were discarded. Of the remaining sample (N=16), participants were better at judging whether the probe matched the cue when the T2 test interval matched the learned interval, L2 (mean [SE] = 0.63 [0.06]), compared to when it did not match the learned interval, L1 (mean [SE] = 0.49 [0.08]; t(15) = −2.69, P = 0.017, Cohen’s d = −0.67). For T1 trials, accuracy did not significantly differ between learning intervals L1 (0.55 [0.08]) and L2 (0.63 [0.07]; t(15) = −1.79, p = 0.09).

### ROI analyses: hippocampus and striatum

For the fMRI analysis, we additionally discarded the data of one participant with corrupted image files, leaving N=15 for further analysis. Activation statistics for each ROI per condition L1xT2 and L2xT2 are listed in **Table 1**. Hippocampal subfields generally showed significantly decreased activity when the temporal interval during retrieval matched the interval during learning (L2xT2). Dorsal striatal structures showed significantly increased activity for both conditions, although activity was higher when the temporal interval during retrieval did not match the interval during learning (L1xT2).

**Table 1.** Region of interest (ROI) results. Signal activation statistics for each ROI during conditions L1xT2 and L2xT2 (degrees of freedom = 14). * significant at false-discovery rate q=0.05; Cd, Cohen’s d; L/R, left/right; Sub, subiculum; DG, dentate gyrus.

| ROIs | L1xT2 |  |  |  |  | L2xT2 |  |  |  |  |
| --- | --- | --- | --- | --- | --- | --- | --- | --- | --- | --- |
|  | M | SEM | T | P | Cd | M | SEM | T | P | Cd |
| LSub | -0.03 | 0.09 | -0.36 | 0.719 | -0.09 | -0.24 | 0.10 | -2.35 | 0.036* | -0.61 |
| LDG/CA4 | -0.15 | 0.11 | -1.36 | 0.193 | -0.35 | -0.34 | 0.09 | -3.73 | 0.002* | -0.96 |
| LCA3/CA2 | -0.10 | 0.08 | -1.22 | 0.241 | -0.32 | -0.32 | 0.08 | -4.24 | 0.002* | -1.10 |
| LCA1 | -0.16 | 0.08 | -2.14 | 0.053 | -0.55 | -0.32 | 0.08 | -3.82 | 0.003* | -0.99 |
| LCaudate | 0.53 | 0.20 | 2.69 | 0.005* | 0.70 | 0.30 | 0.08 | 3.56 | 0.005* | 0.92 |
| LPutamen | 0.25 | 0.15 | 1.69 | 0.101 | 0.44 | -0.08 | 0.09 | -0.86 | 0.398 | -0.22 |
| RSub | -0.05 | 0.06 | -0.95 | 0.37 | -0.25 | -0.21 | 0.07 | -2.96 | 0.01* | -0.77 |
| RDG/CA4 | -0.09 | 0.12 | -0.75 | 0.493 | -0.19 | -0.25 | 0.09 | -2.88 | 0.017* | -0.74 |
| RCA3/CA2 | -0.19 | 0.07 | -2.76 | 0.019 | -0.71 | -0.19 | 0.07 | -2.62 | 0.017* | -0.68 |
| RCA1 | -0.04 | 0.09 | -0.47 | 0.661 | -0.12 | -0.24 | 0.07 | -3.48 | 0.003* | -0.90 |
| RCaudate | 0.50 | 0.19 | 2.68 | 0.004* | 0.69 | 0.29 | 0.10 | 2.95 | 0.012* | 0.76 |
| RPutamen | 0.27 | 0.16 | 1.65 | 0.086 | 0.43 | -0.07 | 0.11 | -0.63 | 0.544 | -0.16 |

Several ROIs showed significant differences in activity between the two conditions. **Figure 3** shows the distribution of regression coefficients between the two conditions for the four hippocampal subfields and the two DS nuclei in each hemisphere. For hippocampal subfields, we found significant deactivations in left CA3/CA2 (t(14) = −3.12, p = 0.008, adjusted- p = 0.035, Cohen’s d = −0.81), left CA1 (t(14) = −2.67, p = 0.013, adjusted-p = 0.035, Cohen’s d = −0.69) and right CA1 (t(14) = −2.91, p = 0.01, adjusted-p = 0.035, Cohen’s d = −0.75). For all other hippocampal subfields, uncorrected p-values were larger than 0.05.

**Figure 3.**
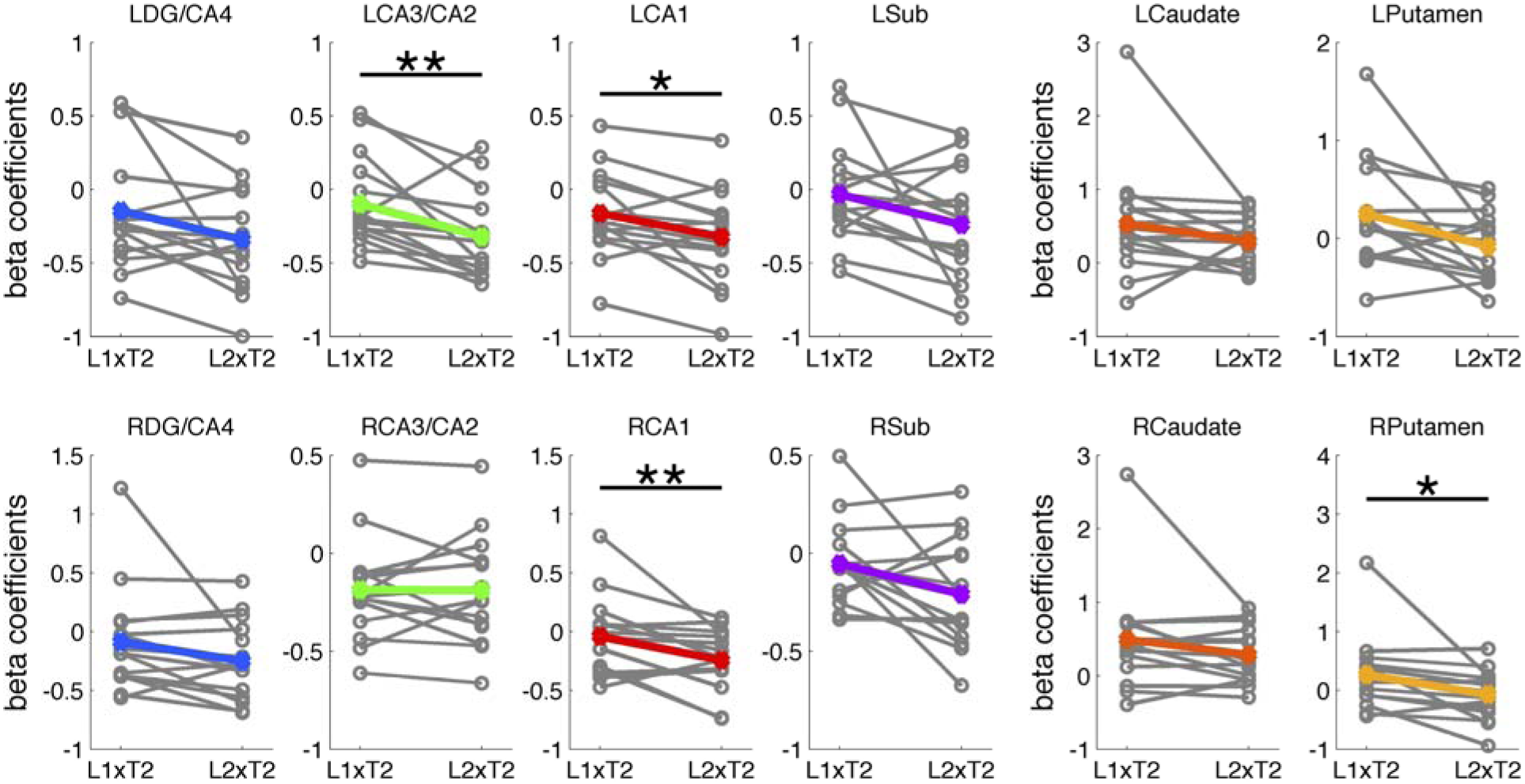
ROI results. Mean activity during conditions L1xT2 and L2xT2 for each of the hippocampal subfields and dorsal striatal nuclei. * P < 0.05, ** P < 0.01.

For the dorsal striatal areas, we found a significant increase in activation in right Putamen (t(14) = −2.46, p = 0.004, adjusted-p = 0.015, Cohen’s d = −0.64). We also found increased activity in left Putamen, but this effect was not significant after multiple comparison correction (t(14) = −2.30, p = 0.031, adjusted-p = 0.063, Cohen’s d = −0.59). Effects in the Caudate were not significant at uncorrected p-values (ps > 0.1).

To further investigate the relation between brain activity and temporal expectation, we correlated activity of left CA1 and right putamen with task performance pooled across both conditions. Given the difference in direction of activity in the two areas, we would expect that better task performance was associated with decreased CA1 activity, but with increased putamen activity. We found a negative correlation (Spearman) between accuracy and CA1 (rho = −0.37, p = 0.043) and a positive correlation between accuracy and right putamen (rho = 0.45, p = 0.012), thereby corroborating the suggestion that these subcortical structures contributed to temporal associative memory performance.

### PPI of the left CA1 × task onto the striatum

In this analysis, we investigated whether functional connectivity between the hippocampal subfield CA1 and striatal nuclei changed with different task conditions. A PPI model with the task contrast of L1xT2 vs. L2xT2 as psychological factor and left CA1 activity as physiological factor was applied to each of the four ROIs of the dorsal striatum. The psychological and physiological vectors were mean-centered and deconvolved prior to calculating the PPI interaction term.

We found a significant psycho-physiological interaction term in left (t(14) = −2.69, p = 0.013, adjusted-p = 0.026, Cohen’s d = −0.70) and right caudate (t(14) = −2.95, p = 0.008, adjusted-p = 0.026, Cohen’s d = −0.76). **Figure 4A** shows the scatterplots of the normalized fMRI signal of the right caudate as a function of fMRI signal in left CA1 for two representative participants. The different regression lines, showing a higher correlation for the L2xT2 condition (black line) compared to the L1xT2 condition (grey line), indicate that the PPI interaction term represented increased hippocampal-striatal connectivity when the tested temporal interval matched the learned interval. The PPI interaction term was not significant for left or right Putamen.

**Figure 4.**
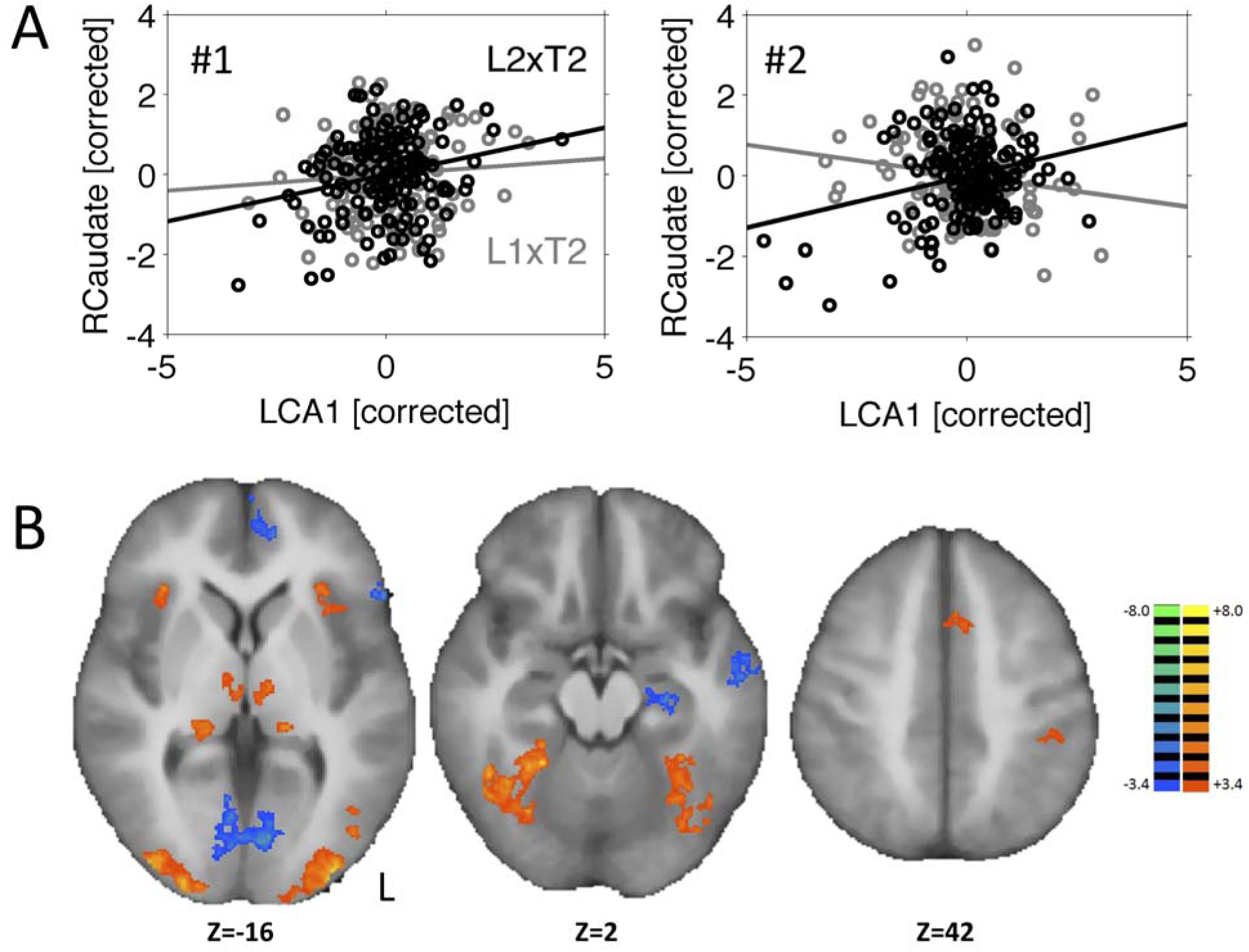
PPI scatterplots and whole-brain results. A) Psycho-physiological interaction (PPI) scatterplots for activity of right caudate as a function of activity in CA1 for L1xT2 (grey) and L2xT2 (black) of two representative participants. B) Areas of significantly (cluster-level corrected) increased (hot colors) or decreased (cool colors) activity for the task, *i.e.*, L1xT2 + L2xT2. Left hemisphere shown on the right in each panel.

The findings of PPI connectivity between CA1 and caudate, but not with putamen, suggested that the two striatal structures may play different but complementary roles in processing of temporal associative context. The task-related change in putamen activity could indicate that it monitors whether the cue-dependent temporal expectancy is met (Mcclure et al., 2003), whereas caudate functional connectivity could indicate an integration of current sensory experiences with (temporal) expectations. In this scenario, caudate should show task-related functional connectivity with putamen. To investigate this possibility, we conducted another PPI analysis of right caudate (having shown the strongest connectivity with CA1) with putamen activity as physiological variable. Results revealed a significant PPI interaction term (t(14) = - 3.10, p = 0.007, Cohen’s d = −0.80). Together, these results indicate that caudate may integrate temporal expectations from (hippocampal) memory with a signal from putamen that indicates whether the expectations are violated.

### Whole-brain analysis

Finally, we conducted a more explorative whole-brain voxel-by-voxel analysis of the main effect of the task, that is, L1xT2 + L2xT2. Voxel-level results were initially thresheld at an uncorrected p value of 0.005 and were further controlled for multiple comparisons at the cluster-level at a false positive rate of 0.05. Results (see **Figure 4B**) showed increased activity during trials of both conditions in bilateral lateral occipital cortex, inferior temporal cortex, supplementary motor area (SMA), left sensorimotor cortex and medial thalamic and lateral geniculate nuclei. Decreased activity was found in left hippocampus, ventral medial prefrontal cortex, posterior parietal cortex and medial occipital cortex at putative primary visual cortex. The contrast of L1xT2 – L2xT2, corrected for multiple comparisons at the cluster-level, revealed no significant effects.

## Discussion

We utilized UHF 7T fMRI to examine how memory-based temporal prediction is represented on the hippocampus and dorsal striatum. Significantly higher activations were detected in several hippocampal subfields, including bilateral CA1 and left CA3/CA2, when temporal context at retrieval did not match the context used during learning (L1xT2), compared to when temporal context did match between learning and testing (L2xT2). The involvement of CA3-CA1 subfields in our task fits with previous neurophysiological studies that showed the involvement of CA subfields in processing temporal associative context. Rat lesion studies have identified CA3 and CA1, but not DG, as critical subfields for the formation of associations between objects separated by a temporal delay (Hunsaker et al., 2006; Hunsaker and Kesner, 2008; Farovik et al., 2010). Moreover, the presence of “time cells” that represent the temporal moments of events have been observed in CA3 (Salz et al., 2016) as well as CA1 (MacDonald et al., 2011, 2013; Kraus et al., 2013). Further, hippocampal activity in humans has also been associated with the encoding of temporal context (Schapiro et al., 2012; DuBrow and Davachi, 2014; Ezzyat and Davachi, 2014; Hsieh et al., 2014; Montchal et al., 2019). Interestingly, some studies showed changes in left hippocampal activity (DuBrow and Davachi, 2014; Ezzyat and Davachi, 2014) while other studies found more activity in the right hippocampus (Schapiro et al., 2012; Hsieh et al., 2014). Perhaps this is due to selective roles of the left and right hippocampus in processing different temporal aspects of memory. Hsieh and colleagues (Hsieh et al., 2014; Ranganath and Hsieh, 2016) proposed that the right hippocampus may be more sensitive to processing the temporal order of items within a sequence, whereas the left hippocampus may process temporal contextual markers such as event boundaries or temporal distance between items. Considering our study, in both learning and testing phases the ordering of cue-target pairs was kept constant while the temporal gap between cue and target varied. Thus, rather than temporal order, our task paradigm manipulated cue-dependent (expected) duration as temporal context, thereby activating left more strongly than right hippocampus.

We also found increased signal amplitude in the right putamen when temporal expectations were violated. Generally, this finding fits with the long-held hypothesis that dorsal striatum is involved in processing of timed stimulus-response gaps (Rao et al., 2001; Coull et al., 2004; Tanaka et al., 2004; Buhusi and Meck, 2005; Wiener et al., 2010; van Rijn et al., 2014), but also extends it to processing of temporal information in associative memory. Particularly, our finding mirrors a previous finding of right putamen activity when temporal expectancy about reward delivery was violated (Mcclure et al., 2003). Thus, in our study, right putamen may have coded for the violation of temporal expectation that arose from associative memory., and fits with the more general notion that dorsal striatum monitors the difference between temporal expectancies and experiences, possibly to optimize future action selection (Seymour et al., 2004).

Importantly, our PPI analysis showed increased functional connectivity between left CA1 and striatum when temporal context during testing matched the learned temporal context. This finding is in line with reports that hippocampus and striatum interact cooperatively in the formation of associative memories (Scimeca and Badre, 2012; Mattfeld and Stark, 2015). Previous studies have shown that the two subcortical structures interact during spatial navigation of learning of relevant locations in space (Voermans et al., 2004; Gengler et al., 2005; Igloi et al., 2010; Woolley et al., 2015). Our findings extend this notion to learning of temporal associations. Moreover, our findings suggest that caudate and putamen may have differential roles in processing of temporal associations. The task-related change in activity suggests that putamen monitors whether temporal expectations are met, while the task-related change in connectivity with CA1 suggests that caudate integrates sensory experiences with memory-based expectations that arise from associative memory. This reasoning fits with previous results showing that putamen, but not caudate, is activated when expectations about temporal delays (Mcclure et al., 2003) or expected (non-temporal) outcomes are not met (Brovelli et al., 2011). Concurrently, caudate nuclei may integrate information about temporal expectancy and actual experience in order to optimize the selection of when and how to respond.

Some final remarks about our study are warranted. The sample size, although arguably small, is comparable with other recent 7T imaging studies (*e.g.*, (Raemaekers et al., 2013; Ten Oever et al., 2016)). We optimized statistical power by restricting the analysis search space to a small set of task conditions and subcortical regions-of-interest. A possible limitation of the study, however, is that general performance accuracy was lower than that in a previous study, in which participants were not scanned (van de Ven et al., 2017). This difference in performance suggests that the scanner environment could have affected task performance.

In conclusion, our fMRI study demonstrated that elapsed time in associative memory could function as an important mnemonic context for the hippocampus and striatum at retrieval. Our findings revealed CA3-CA1 hippocampal subfields and dorsal striatum as important neural correlates involved in processing temporal information of memory. Moreover, these regions were functionally connected when the temporal context at retrieval was identical to the learned temporal context. The results extend current knowledge of memory and time beyond hippocampal areas and start to explain how contextual information engrained in memory is perceived and analyzed in the human brain.

## Acknowledgments

This study was partially funded by an MBIC-FPN (Maastricht Brain Imaging Center (MBIC) and Faculty of Psychology and Neuroscience (FPN)) grant. All authors report no conflict of interest. The data that support the findings of this study are available from the corresponding author upon reasonable request <citation provided when accepted>.

